# Efficient Precision Dosing Under Estimated Uncertainties via Koopman Expectations of Bayesian Posteriors with Pumas

**DOI:** 10.1101/2021.01.25.428134

**Authors:** Chris Rackauckas, Vaibhav Dixit, Adam R. Gerlach, Vijay Ivaturi

**Affiliations:** Massachusetts Institute of Technology and Pumas-AI Inc.; Pumas-AI Inc.; United States Air Force Research Laboratory; University of Maryland, Baltimore, School of Pharmacy and Pumas-AI Inc.

## Abstract

Personalized precision dosing is about mathematically determining effective dosing strategies that optimize the probability of containing a patient’s outcome within a therapeutic window. However, the common Monte Carlo approach for generating patient statistics is computationally expensive because thousands of simulations need to be computed. In this manuscript we describe a new method which utilizes the Koopman operator to perform a direct computation of expected patient outcomes with respect to quantified uncertainties of Bayesian posteriors in a nonlinear mixed effect model framework. We detail how quantities such as the probability of being within the therapeutic window can be calculated with a choice of loss function on the Koopman expectation. We demonstrate a high performance parallelized implementation of this methodology in Pumas^®^ and showcase the ability to accelerate the computation of these expectations by 50x-200x over Monte Carlo. We showcase how dosing can be optimized with respect to probabilistic statements respecting variable uncertainties using the Koopman operator. We end by demonstrating an end-to-end workflow, from estimating variable uncertainties via Bayesian estimation to optimizing a dose with respect to parametric uncertainty.

## 1 Introduction: The Importance of Precision Dosing

Quantitative clinical pharmacology model-based dose selection in the wider context of model-informed drug development is widely used by the pharmaceutical industry. However, the applications of such technologies have had little to no impact in the clinic till about a few years ago. The challenge for model-based dosing lies in the idea that drug development programs are often incentivized to devise dosing recommendations in the product label primarily for ease of prescribing. This extends to the dosing guidelines for special populations where the label specifies dosing based on certain cut-points for prognostic factors such as age, weight, kidney function or disease status. Historically, the primary reason for such simplified label recommendations was to facilitate approval for the average patient. This, however, is no longer the case as advances in quantitative clinical sciences (*e.g.,* pharmacometrics) and technology enable an individualized approach to drug therapy. For much therapeutics, this signifies an important paradigm shift from a predefined dose to a more tailored and personalized dose aimed to increase efficacy and reduce toxicity.

For effective care, dosing for each patient can be individualized to achieve some desired target goal, such as serum concentrations, or an effect such as bacterial kill or longitudinal glucose measurements over time. Frequent observations of these targets at optimal times will facilitate dose adjustment as needed. However, it is also important that these adjustments are made in real time to avoid delays in decision making that can lead to unintended consequences with regards to safety or loss of efficacy. Target goals are usually defined by therapeutic windows, especially for drugs with a narrow safety margin. For such drugs, therapeutic drug monitoring (TDM) is used to measure *e.g.,* serum concentrations frequently. Based on the observed concentrations and prior clinical experience these drugs are then classified into sub-therapeutic, therapeutic 1 or toxic levels. Clinical and therapeutic committees in hospital management meet and develop guidelines as to what the acceptable ranges of drug concentration are, which are subsequently used by clinicians to adjust doses to achieve those ranges. Over the years, therapeutic windows have been defined for multiple drugs. While the list of narrow therapeutic index differs across regulatory bodies, a comprehensive list of narrow therapeutic index drugs can be found in DrugBank Online Database^*^. Precision dosing has seen a widespread implementation in the areas of infectious disease and transplant, while more recently application in oncology and epilepsy have also surfaced [9].

However, adequately modeling the probability that a drug concentration remains within the therapeutic window remains a challenge. Monte Carlo methods compute the probability by simply computing millions of samples and checking the percentage of results which lie within in the window. Given that model simulations can be computationally time consuming and have slow convergence, it can be impractical to achieve accurate probabilistic estimates through this classical approach.

To address this issue, we describe a mathematical formalism and a new method to facilitate precision dosing. We utilize the Koopman expectation for computing expectations and probabilities of model quantities with respect to estimated parametric uncertainties. We show how to phrase the probability of internal drug concentrations being within the therapeutic window as a quantity that can be calculated via the Koopman expectation. We demonstrate how the Koopman expectation allows for 50x-200x more efficient estimation of these probabilistic quantities compared to Monte Carlo which enables optimization of dosing schemes with respect to uncertainty in less than a second. Additionally, we showcase how the quadrature approach to the calculation of the Koopman expectation yields informative error bounds on the probabilistic estimates, improving their ability to be used and trusted in practice. Together this demonstrates a practical approach for real-time decision making under uncertainty that can be applied to precision dosing scenarios.

## 2 Background: Bayesian Estimation of Nonlinear Mixed Effects Models

Dynamics of drug concentrations (pharmacokinetics, PK) and patient outcomes (pharmacodynamics, PD) on patient populations are commonly modeled via nonlinear mixed effects models (NLME) [4]. The dynamics of the PK/PD system are defined in terms of an ordinary differential equation

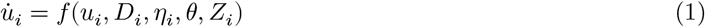

where *D_i_* is the dosage regimen, *θ* are the fixed effects, *η*_*i*_ are the random effects for patient *i* and *Z*_*i*_ are the covariates of patient *i*, known measurable quantities of a patient such as their age, weight, or sex. For this model, we assume that *η*_*i*_ ~ *N*(0, Ω), and thus the expected patient outcome given only the prior known information *Z*_*i*_ is the dynamics predicted by *θ* with *η*_*i*_ = 0.

Given observations *d* of the underlying dynamical system *u*, fixed effects *θ* and random effects variance Ω, the population likelihood *L* is given by

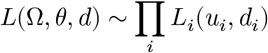

where *L*_*i*_ is the likelihood for individual *i* which is a chosen functional form. The likelihood is the probability of seeing the data *d*_*i*_ given a distribution around the predicted value. For example, a common choice of *L*_*i*_ is the normal distribution centered around the subject’s predictions, *i.e., L*_*i*_ ~ *N*(*u*_*i*_, *σ*) for some fixed variance *σ*. Note that with this choice of likelihood, choosing *θ* and *η*_*i*_ which maximize *L*_*i*_ is equivalent to choosing the parameters which minimize ||*u*_*i*_ − *d*_*i*_||.

Since we wish to capture the uncertainty in the parameters, instead of performing a deterministic maximum likelihood procedure, we resort to a Bayesian estimation of the posterior distributions for *θ* and *η*_*i*_ given assumed prior distributions [3, 22, 24, 18, 27]. Common probabilistic programming languages, such as Stan [7] or Turing [10], can be utilized to determine these posterior distributions using Markov Chain Monte Carlo (MCMC) techniques such as Hamiltonian Monte Carlo [1, 2, 15]. We will demonstrate how estimated distributions of NLME parameters computed using the Pumas^®^ software [20] can be reinterpreted as probability distributions via a kernel density estimate (KDE) [16] and used within a Koopman expectation framework to greatly accelerate probabilistic estimates and optimization of clinical choices with respect to uncertainty.

## 3 Uncertainty-Aware Personalized Precision Dosing via the Koopman Expectation

### 3.1 The Frobenius-Perron and Koopman Operators

The mathematical background of the problem and the Koopman expectation is explained in detail in [11]. This is adapted here specifically for pharmacology applications. Assume that Bayesian estimation has given sufficient estimates of posterior distributions for *θ* and *η*_*i*_ and we wish to answer the following question: what is the optimal dosage regimen *D_i_* for patient *i*? For the rest of this manuscript we will be focusing on this single patient *i* and thus the subscript will be dropped.

To motivate the application of the Koopman operator for choosing the optimal dose, we first introduce the Frobenius-Perron (FP) operator on dynamical uncertainty distributions [17]. Define *S*(*x*) = *u*(*T*) where *u*(0) = *x*. In other words, *S*(*x*) is the solution of the dynamical system where the initial condition is *x*. We wish to compute probabilistic statements on patient outcomes with respect to uncertainty of the initial condition. First, we note how uncertainties in parameters (*η* and *θ*) are incorporated into this calculation. For the ordinary differential equation, we can use the extended system:

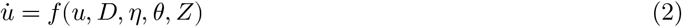

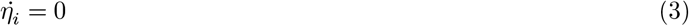

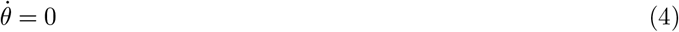

where *η*(0) = *η* and *θ*(0) = *θ*, and thus the parametric uncertainties can be treated as uncertainties in the initial condition on this extended system. Thus without loss of generality, we model the uncertainty via the initial conditions in a probability space 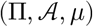 where 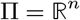. For a given dynamical system 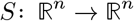, its associated FP operator, *P*_*S*_, is defined such that the following equality is satisfied:

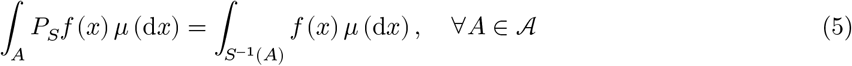

where *S*^−1^ (*A*) is the counter-image of *A* and *f* is the posterior distribution given by the Bayesian estimation [17]. This equivalence is depicted in Figure 1. In terms of precision dosing, the uncertainty in the fixed and random effects, *i.e., θ* and *η* according to the probability distribution *f*(*x*), induces a probability distribution on the solution of the dynamics *u*(*t*). This pushforward of the probability distribution through the dynamical system is the FP operator, *i.e.,* 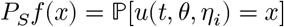.

**Figure 1:**
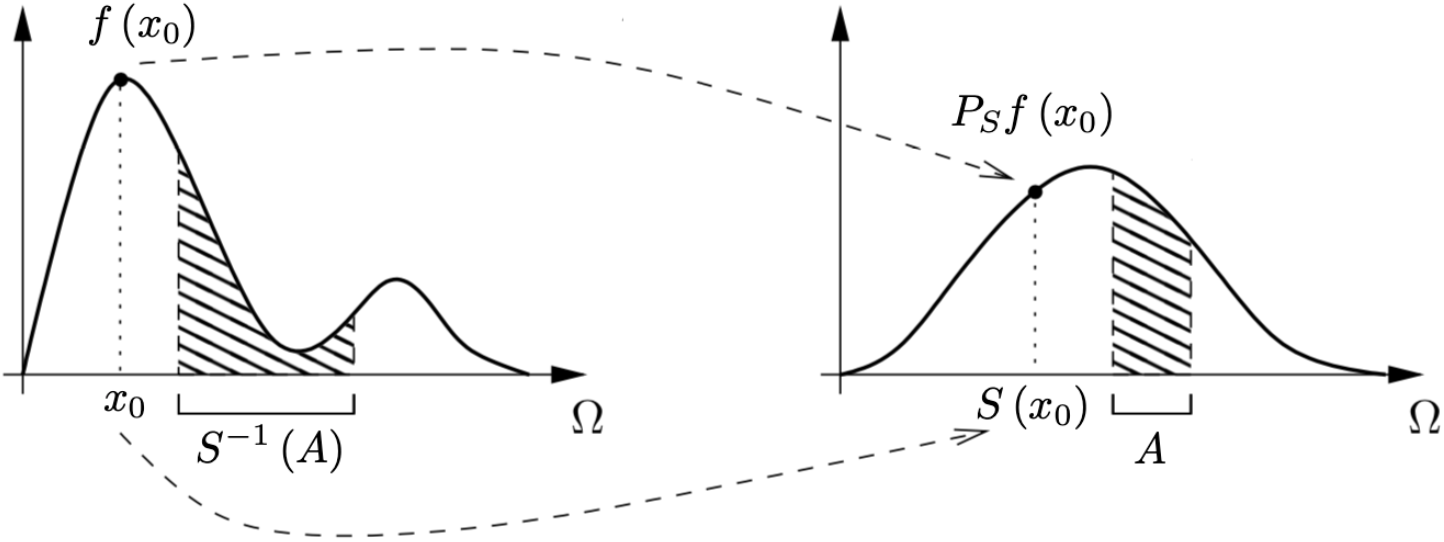
Graphical representation of Eq. 5, where shaded regions have equal area. (Figure adapted from [26])

If *S* is both measurable and nonsingular, then *P*_*S*_ is uniquely defined by Eq. 5 [17]. Monte Carlo estimation of probabilistic quantities of the dynamical system’s solution thus correspond to approximating the pushed forward probability mass *P*_*S*_*f* by direct sampling of initial conditions from the probability distribution *f*.

In contrast, the Koopman operator *U*_*S*_ is defined as

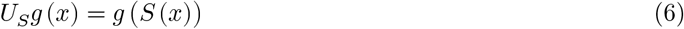

where *g*(*x*) denotes the observed quantities of the solution to the dynamical system (which we will later call the cost function). Note that *U*_*S*_ is adjoint to the FP operator, *P*_*S*_, *i.e.,*

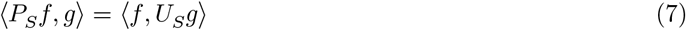

where 〈·, ·〉 is the inner product. Eq. 7 can be rewritten as

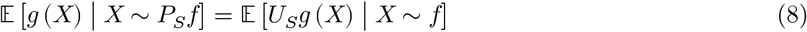

We refer to the left- and right-hand sides of Eq. 8 as the *Frobenius-Perron* (FP) and *Koopman Expectations*, respectively. Figure 2 provides a 1D illustration of Eq. 8. The top row represents the FP Expectation while the bottom row represents the Koopman Expectation. For the top row, the PDF *f* (dashed line) is pushed to the right through the system dynamics via *P*_*S*_ and an inner product is taken with *g* (solid line). The expected value, 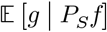, is represented by the area of the shaded region. Conversely, on the bottom row, the function *g* is pulled to the left through the system dynamics via *U*_*S*_. The expected value, 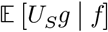, is represented by the area of the shaded region. The areas of the two shaded regions are equal.

**Figure 2:**
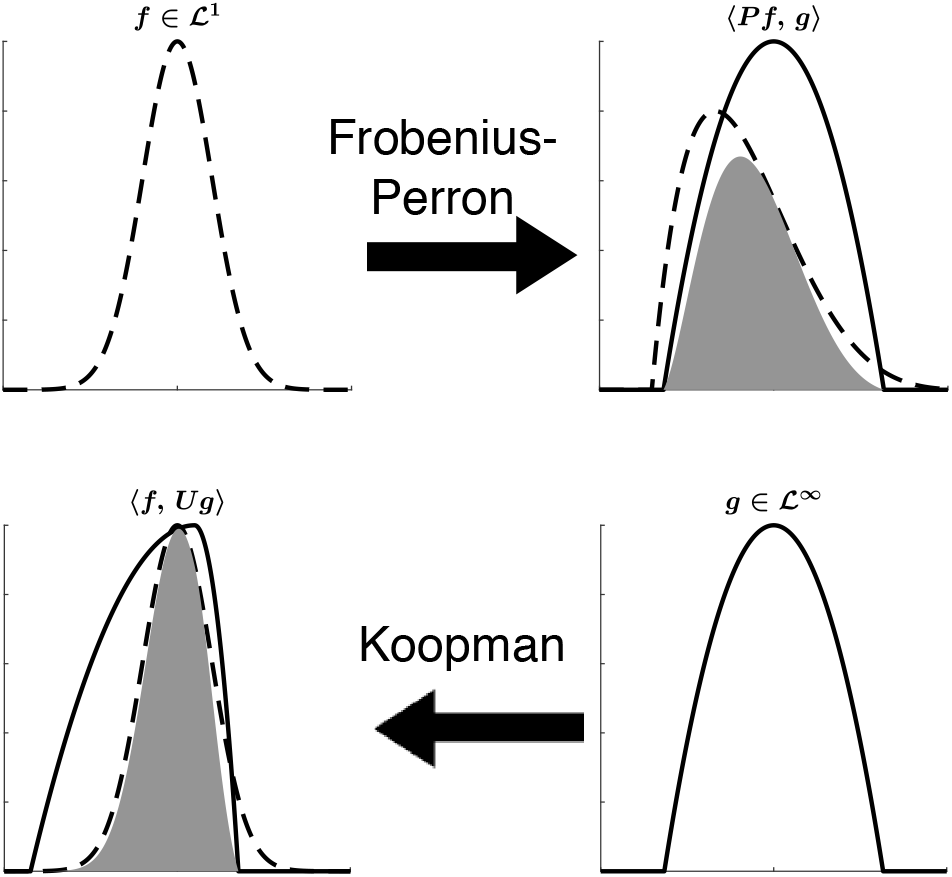
Illustration of the FP and Koopman operator adjoint property. The inner products, represented by the area of the filled regions, are equivalent. (Figure reproduced from [19])

### 3.2 Precision Dosing as Optimization Under Uncertainty

Precision dosing under uncertainty can be phrased as a problem of optimal decision making under uncertainty. Concretely, the optimal dose is one which has the highest expectation of good patient outcomes, *e.g.,* being within a therapeutic window. In other words, we wish to find the dosing schedule *D* which optimizes the expectation with respect to a cost function on the solution of the dynamical system. If we let *g*(*u*(*T*)) be the cost associated with a given patient outcome, this corresponds to finding *D*\* such that

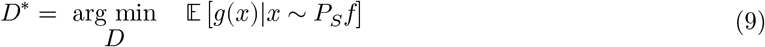

Thus the common way to perform dosing optimization with respect to parametric uncertainty is to utilize a Monte Carlo estimation of *P*_*S*_*f* in order to evaluate the expectation, *i.e.,*

1. Sample parameters *θ* and *η*_*i*_ from the uncertainty distribution *f*
2. Solve the dynamical system to compute *S*(*x*) for each set of parameters
3. Compute *g*(*S*(*x*)) on each set of parameters, and take the discrete average

This procedure is computationally expensive since it requires the solution of many differential equations. However, using the relationship of Equation 8, we see that the argument of the optimization problem is simply the Frobenius-Perron expectation, and thus it can equivalently be rephrased in terms of the Koopman expectation [11]:

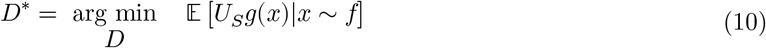

Notice from Figure 2 that the Koopman operator pulls back to the original probability distribution. Thus we can explicitly represent the calculation as:

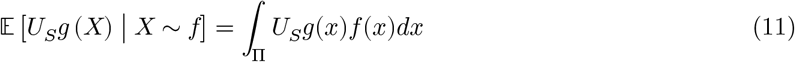

In other words, this means the desired expectation can be calculated via a multidimensional quadrature where *U*_*S*_*g*(*x*) is the solution of the dynamical system at parameters determined by the quadrature procedure and *f*(*x*) is the evaluation of the probability distribution at the quadrature points.

### 3.3 Representation of Therapeutic Windows in Cost Functions

Clinical experience via TDM establish safety guidelines known as the therapeutic window. For example, for a given drug the area under the curve (the total concentration or AUC) of the drug over 24 hour periods may be known to be safe when it is between 200 and 400 mg.hr/L. Given variability in drug disposition and elimination, the goal is to optimize the probability that the dose will be in the therapeutic window with respect to the uncertainty in the patient-specific effects.

To calculate this probability, let *g*(*X*) = *χW*(*X*) where *χW* is an indicator function for the therapeutic window *W*, *i.e., g* is a function defined as 1 if the solution to the differential equation with parameters (*θ*, *η*, *D*, *Z*) falls in the therapeutic window, and 0 otherwise. With this definition of the cost function, we note that the expectation of a characteristic function is the probability that the event occurs, and as such this definition of *g* gives rise to:

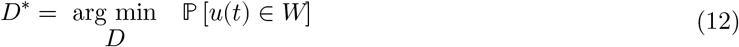

Therefore under this choice of *g* the optimal dosing regimen *D*\* computed by the optimization over the Koopman expectation is the dosing regimen that has the maximal probability of the patient’s outcomes to be in the therapeutic window. We note in passing that this formulation is slightly more general, as it allows for a quantification of “badness”, *i.e.,* the cost could be a distance from the therapeutic window which encapsulates the idea that being close to the therapeutic window might be sufficient while being far away incurs a greater risk on the patient. Generalizations to this computation which include higher order statistics and process noise are given in [11].

## 4 Efficient Computation of the Koopman Expectation in Pumas^®^

### 4.1 Description of the Implementation and Features

To demonstrate the utility of this method for performing dosage optimization under uncertainty, we created a high-performance implementation of the Koopman expectation in Pumas^®^. Equation 11 was implemented by utilizing DifferentialEquations.jl [21] to calculate *U*_*S*_*g* given a Pumas^®^ [20] specification of a dynamical system and the multidimensional integral was calculated using Quadrature.jl, a wrapper library over common quadrature methods such as Cuba [13] and Cubature [12, 6]. This integration implementation allows for a batch solve that parallelizes the computation over the quadrature points, allowing for multithreaded, distributed, and GPU acceleration of the quadrature. The quadrature techniques were made differentiable in order to allow automatic differentiation use for optimization under uncertainty.

### 4.2 Calculating Probabilities With Respect to Theophylline Dosing

The following code demonstrates the use of the Koopman expectation for the calculation of the probability that the AUC will be below 300 on the Theopylline model. The nonlinear mixed effect model’s definition is defined using Pumas^®^’ standard @modelmacro form:

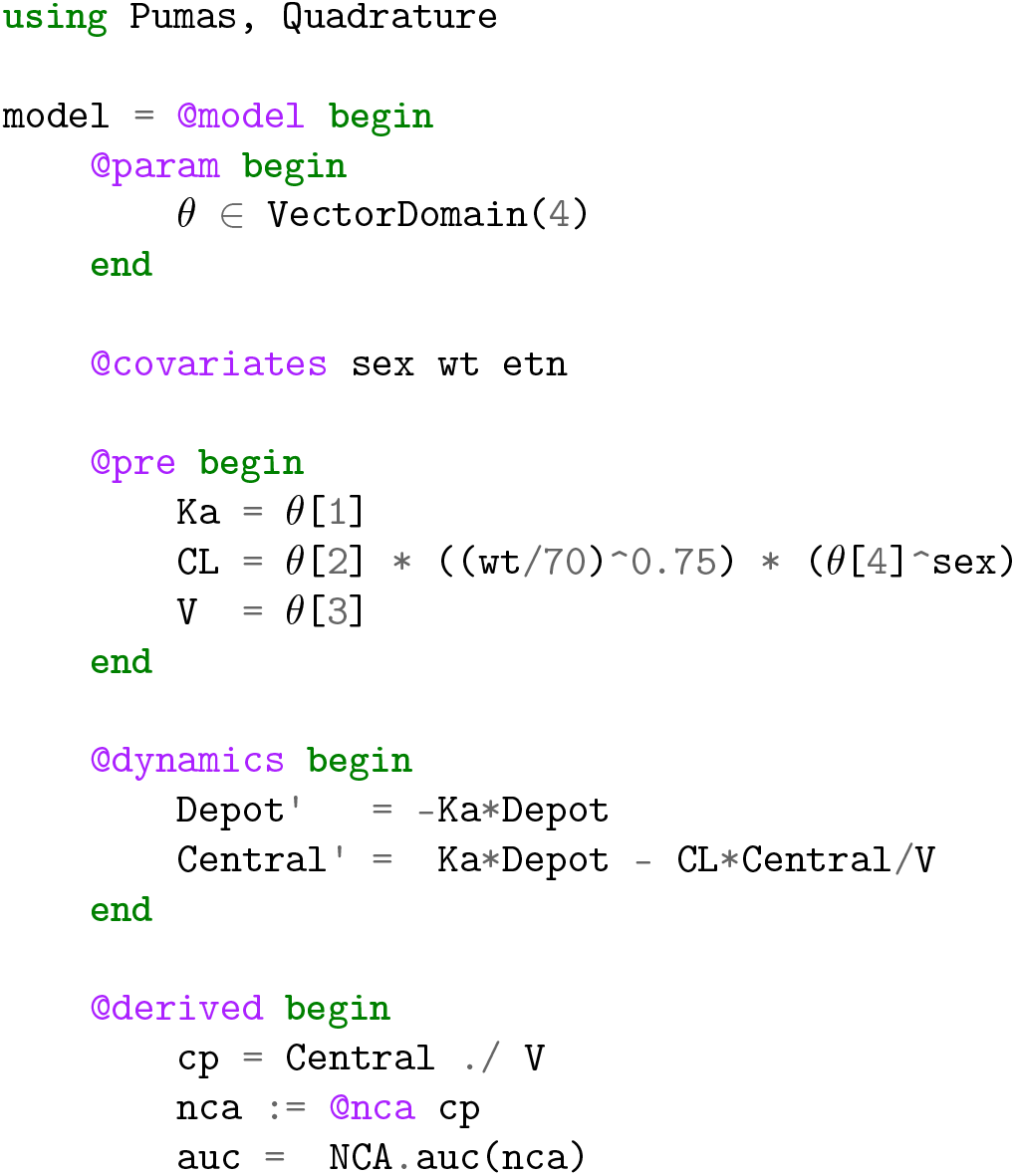

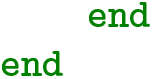

Then to define the subject and the dosage regimen, we read in a dataset:

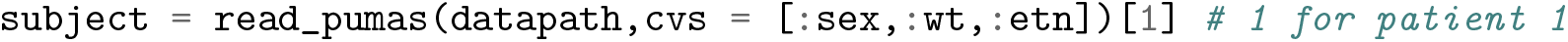

Next we input the parameter uncertainty distributions. This would normally come from Bayesian posteriors as described in Section 2, but here we will directly prescribe distributions as an illustrative example.

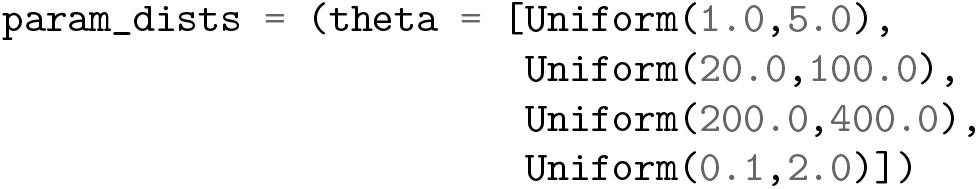

We then define the therapeutic window via the cost function *g* on the observables. Notice that we used the Pumas^®^ NLME integration with NCA to compute the AUC as part of the derived variables from the model, and thus the AUC exists as one of the observed outputs of the system. We can thus define the therapeutic window via:

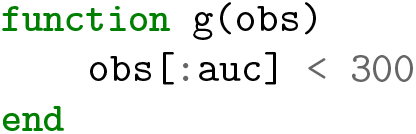

Now we simply call expectation and tell it to use the KoopmanExpectation method. In there we designate that we would like to use the HCubatureJL quadrature method, which tends to be efficient for low dimensional integrands (<8 uncertain variables).

~~~
expectation(g,KoopmanExpectation(CubatureJLp()),model,subject,param_dists)
~~~

This computation will then compute the given expectation to the desired tolerance. Given its differentiability, this function can then be used inside of an optimization loop with defined dosage regimens to thus perform dosage optimization under uncertainty.

### 4.3 Efficiency and Robustness of the Koopman Expectation vs Monte Carlo

This extra formalism is useful because of the tremendous performance and feature benefits. To demonstrate this, we compared the KoopmanExpectation to the MonteCarloExpectation option with various increasing choices of imaxiters (allowed number of ODE solves) to determine the rate of convergence of the two methods for calculating the probability of patient outcomes occuring in the therapeutic window. Figure 3 demonstrates two results. One is that the Koopman method converges to give stable probability estimates of the second digit with approximately 50× less ODE solves. Secondly, by utilizing the HCubature method the Koopman calculation not only determines the probability but also gives numerical error bounds on the probability estimate, allowing the user to effectively know the uncertainty introduced by the numerical error. Such a bound is highly difficult to calculate with Monte Carlo estimates given the slow rate of convergence of the variance. Together, this demonstrates the Koopman expectation as both a method for efficiency and robustness.

**Figure 3:**
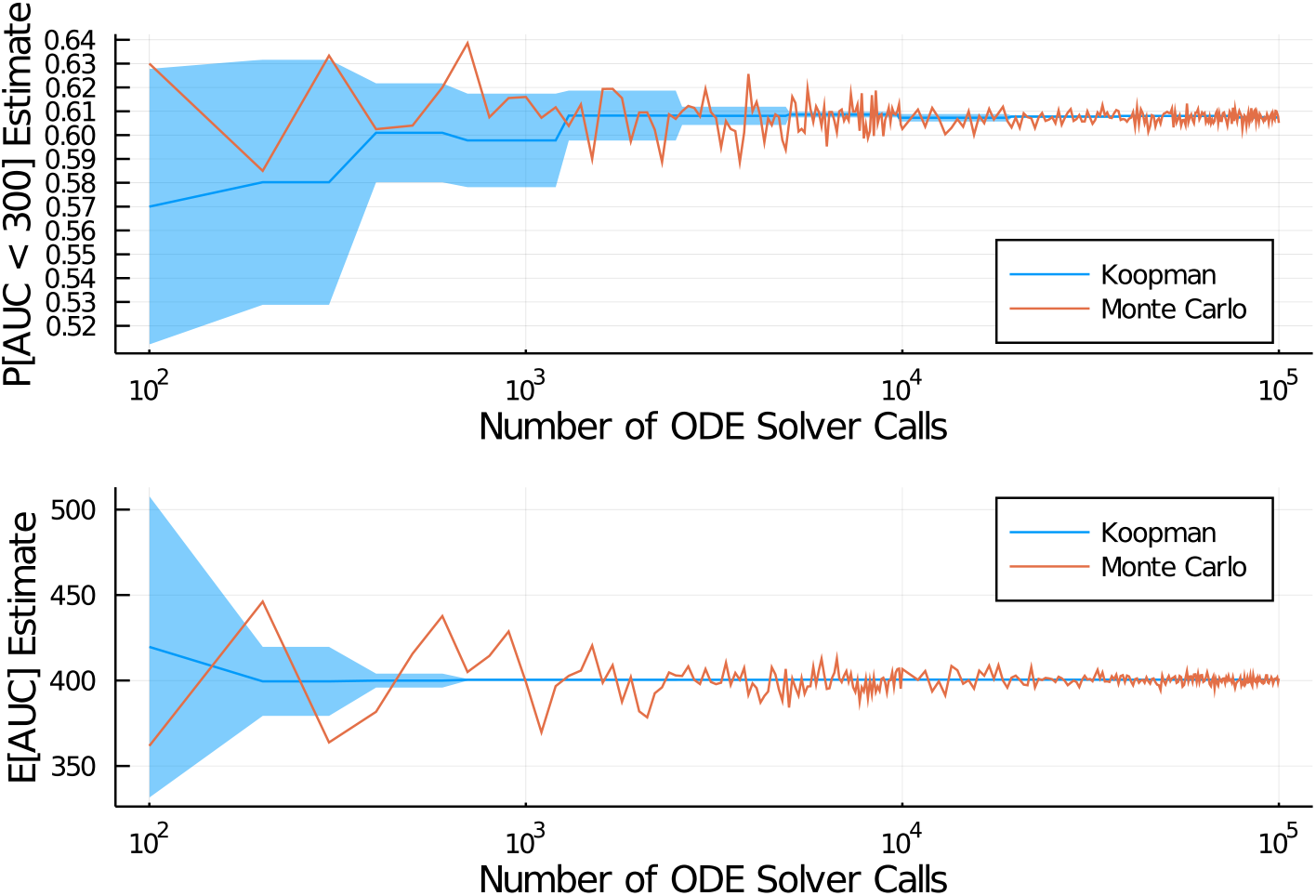
Convergence of the Koopman Expectation calculation via Cubature’s CubatureJLp on the integral (blue line) vs Monte Carlo (orange line), measured in terms of the number of ODE solver calls required. Note that the cubature integration method used with the Koopman Expectation comes with a free error estimation (blue region). **Top:** *P* [*AUS* < 300]. **Bottom:** *E*[*AUS*].

While the quadrature-based Koopman significantly outperforms Monte Carlo, we note that the chosen observable, a characteristic function, was a discontinuous function. The efficiency of quadrature methods can be considerably improved when the integrands are smooth. Thus to test the efficiency in the smooth case, we used the Koopman method to estimate the expected value of the AUC via:

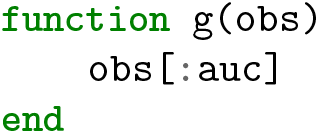

In this case Figure 3 shows that it takes only 500 ODEs solves for the Koopman method to certify that *E*[*AUS*] ≈ 400 ± 1. Meanwhile the Monte Carlo approach is unable to establish a stable estimate within 1 unit with 100,000 ODE solves, with the estimate at 99500 solves giving 398.91 and 10000 giving 401.13. Therefore we see that the Koopman method in this case is effectively 200× as efficient as the classical Monte Carlo method.

### 4.4 Accelerated Bayesian Precision Dosing Using Pumas^®^

In this next example we wish to use the accelerated expectation estimate to optimize a dosing scheme with respect to a probabilistic quantity. We will do so on the Theopylline model with the included example data in Pumas^®^. Let’s start by defining a one-compartment model and fitting it using Bayesian estimation against the example data. This looks like:

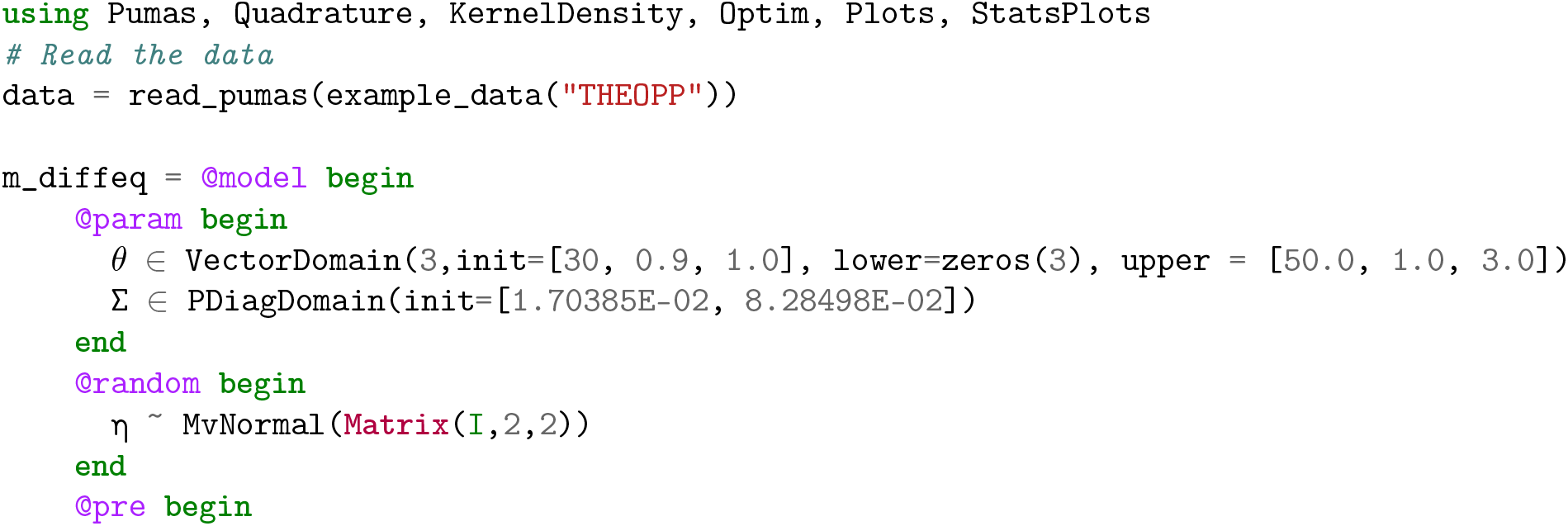

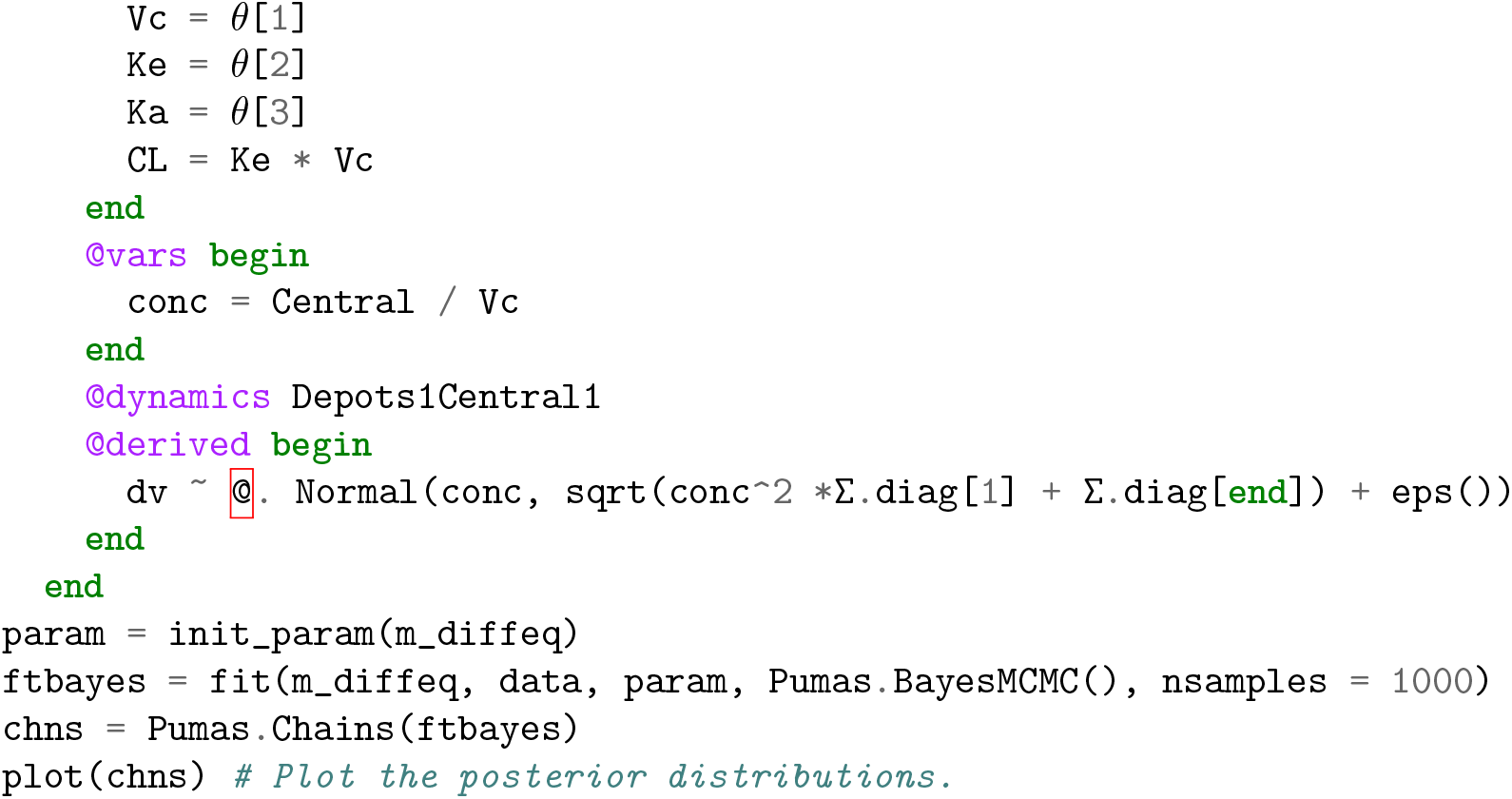

A plot of the posterior distributions for the *θ* terms is given in Figure 4. Given these Bayesian estimates, we can do one of two things to get uncertain parameter estimates. One way is to directly use the Kernel Density Estimate (KDE) to reinterpret the chains into probability distributions. This is done as follows:

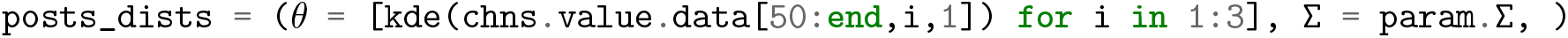

**Figure 4:**
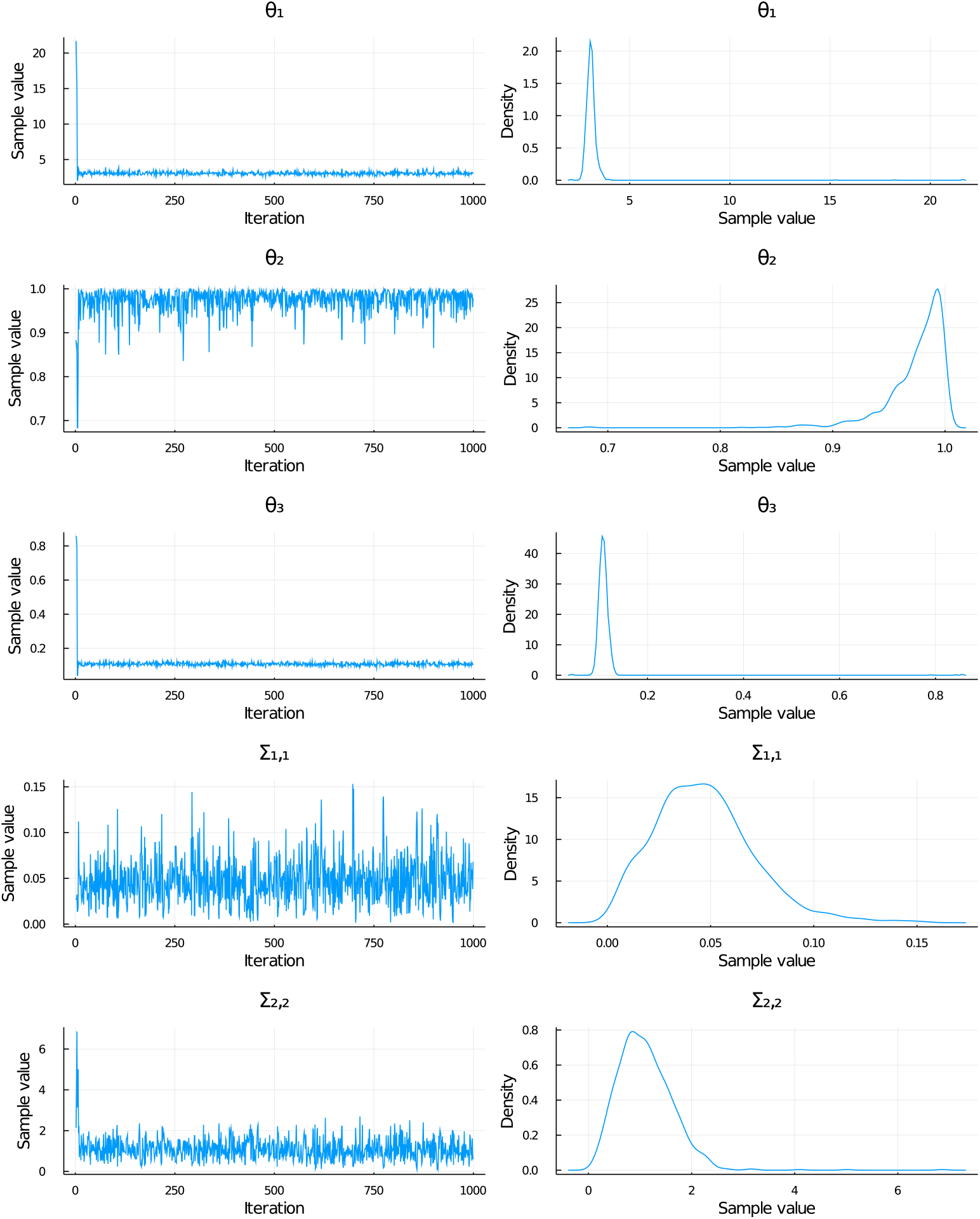
Posterior distributions and MCMC Chains obtained using HMC in Pumas^®^ for bayesian estimation of parameters

This is exactly the same as how the density is shown in Figure 4. However, one can also simplify the distributions by using a maximum likelihood fit against a standard distribution. For example, we can find the best fitting normal distributions for our parameters using the fit overloads from Distributions.jl as follows:

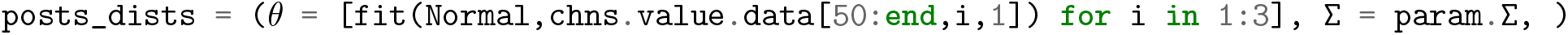

We will continue the tutorial using these distributions for simplicity, though either will give similar results. Note that to stabilize the distributions we chopped off the earliest part of the chain, a procedure known as burn in to reduce the effect of the chain’s random starting guess [8, 14, 25, 5, 23].

Now let’s find the starter dose for which, on average given the estimated uncertainty in the population parameters, has the best chance of having an AUC around 30 in the first five hours. To do so, we make our observable be the AUC calculation using the NCA.jl module included in Pumas^®^:

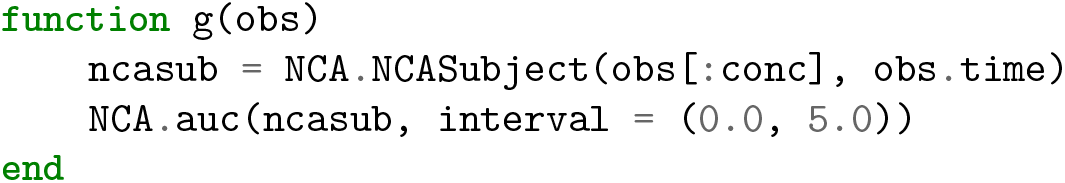

Next we design a new model which makes the maximum and minimum allowed values for the parameters the same as the maximum and minimum of the estimated Bayesian posteriors (to improve the Koopman calculation) and use Optim.jl to find the dose whose average AUC is 30.

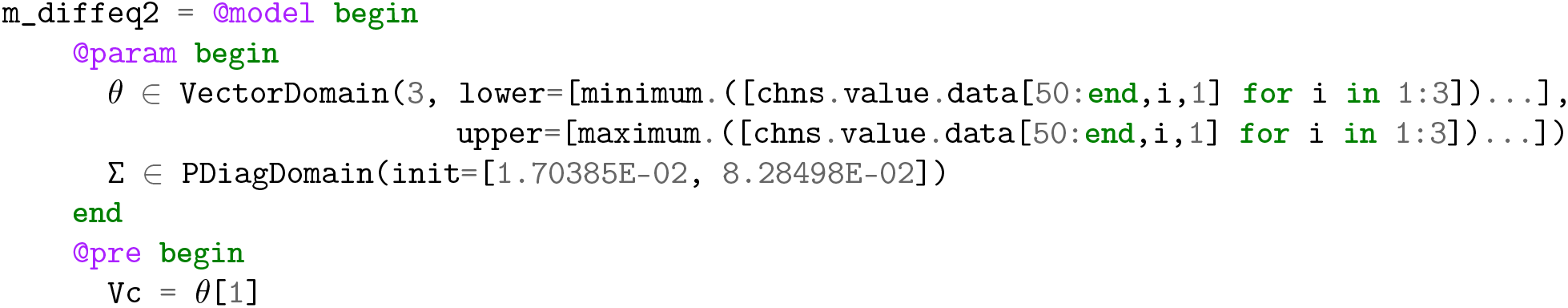

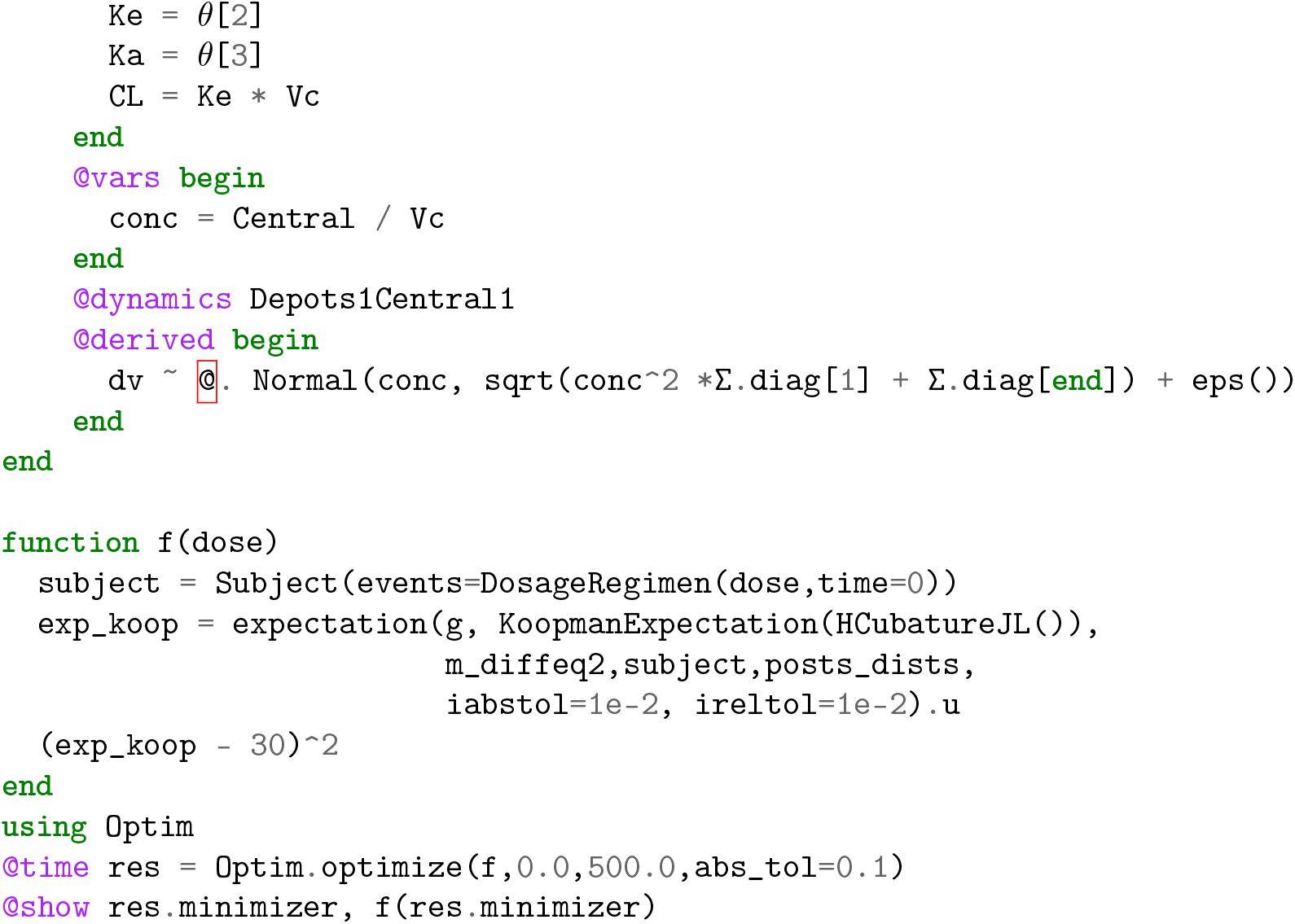

The optimization suggests that the optimal dose to choose to achieve an AUC of 30, respecting the long uncertainty tails in the parameter estimates, is approximately 311. The total time for the optimization was approximately 0.05 seconds.

For reference we can consider the time against Monte Carlo. Accurate gradients are required in order for the optimization to be stable, and thus the optimization procedure needs more digits of accuracy that required by the user. Thus an optimization of the AUC to two digits requires at least 3 stable digits. We find that needs approximately 50000 ODE solves via Monte Carlo, via the code:

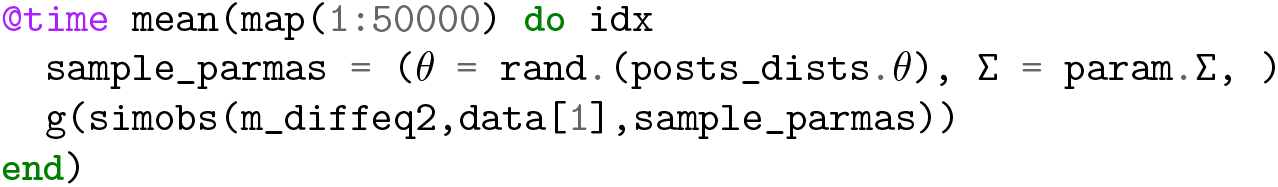

which in three runs produced [36.41,36.45,36.47], showing the 3rd digit is almost stable. However, the time to calculate this estimate is approximately 1.1 seconds, meaning **the entire uncertain dosage optimization with the Koopman expectation is faster than accurately calculating the expectation using Monte Carlo at one point!** We can then see the total effect when trying to optimize a dose with respect to uncertainty:

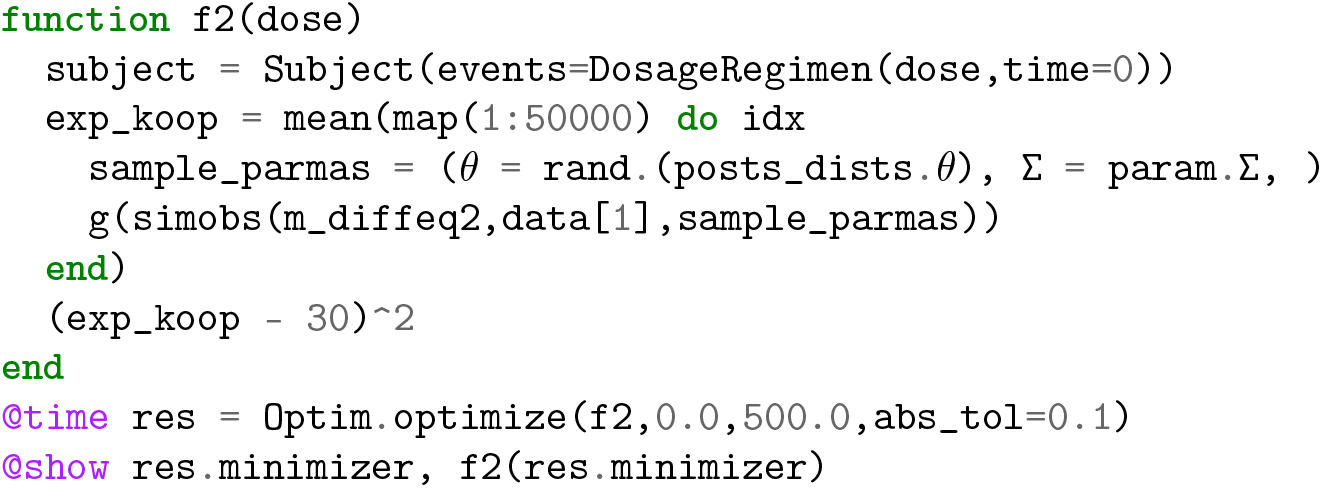

This finds the optimal dose to be around 302 before the optimizer exits due to the stochastic variation of the objective function (reaching a final loss of approximately 41, as opposed to the previous < 1*e* − 8). Multiple runs of the optimizer may be required as the stochastic objective can cause the optimizer to have non-deterministic behavior. In addition, it took approximately 8.5 seconds, or around 170× as long as the Koopman method while not attaining the same accuracy. Because this is done purely by reducing the number of ODE solves required to calculate the expectations, it is expected that such gaps continue to exist as models and data get more expensive to simulate and estimate.

## 5 Discussion

As personalized precision dosing becomes more commonplace, efficient and robust methods for optimizing dosing with respect to probabilistic outcomes will become more central to pharmacometric practice. As personalized computations become pervasive, more accurate models will be required, and the computations will need to migrate to the patient, living on embedded or mobile devices with little compute power. In this manuscript we demonstrated how utilizing the Koopman expectation can accelerate the computation 50×-200× over Monte Carlo while giving numerical error bounds for improved clarity of the results. This methodology is implemented with the high performance differential equation solvers of DifferentialEquations.jl within the Pumas^®^ pharmacometrics suite so that existing models can automatically be compatible with the accelerated computation. Being a pure Julia software stack, this methodology can compile to mobile devices like ARM for direct deployment to patients. Subsequent studies will detail the effect of this methodology in real-world scenarios through its effect on the Lyv (www.lyv.ai) dosing system.

## Footnotes

* https://go.drugbank.com/categories/DBCAT0039722

## Notes

### Competing Interest Statement

C.R., V.D, and V.I. are affiliated with Pumas-AI.

